# Along-tract quantification of resting-state BOLD hemodynamic response functions in white matter

**DOI:** 10.1101/2022.06.09.495555

**Authors:** Kurt G Schilling, Muwei Li, Francois Rheault, Zhaohua Ding, Adam W Anderson, Hakmook Kang, Bennett A Landman, John C Gore

## Abstract

Detailed knowledge of the BOLD hemodynamic response function (HRF) is crucial for accurate analysis and interpretation of functional MRI data. Considerable efforts have been made to characterize the HRF in gray matter (GM) but much less is known about BOLD effects in white matter (WM). However, recent reports have demonstrated reliable detection and analyses of WM BOLD signals after stimulation and in a resting state. WM and GM differ in energy requirements and blood flow, so neurovascular couplings may well be different. We aimed to derive a comprehensive characterization of the HRF in WM across a population, including accurate measurements of its shape and its variation along and between WM pathways, using resting-state fMRI acquisitions. Our results show that the HRF is significantly different between WM and GM. Features of the HRF, such as a prominent initial dip, show strong relationships with features of the tissue microstructure derived from diffusion imaging, and these relationships differ between WM and GM, consistent with BOLD signal fluctuations reflecting different energy demands and differences in neurovascular coupling between tissues of different composition. We also show that the HRF varies significantly along WM pathways, and is different between different WM pathways. Thus, much like in GM, changes in flow and/or oxygenation are different for different parts of the WM. These features of the HRF in WM are especially relevant for interpretation of the biophysical basis of BOLD effects in WM.

## Introduction

Functional MRI (fMRI) based on blood oxygenation level dependent (BOLD) contrast is well-established as a technique to map cortical activity in the brain. BOLD signals indirectly report neural activity, and are characterized by a hemodynamic response function (HRF) which describes the effects of transient changes in blood flow, volume and/or oxygenation (1, 2). The HRF has been shown to vary in amplitude, timing, and shape across brain regions, cognitive states, with ageing, and pathology (3-5), yet accurate estimates of the HRF in any location are crucial for analysis and interpretation of fMRI data.

To date, most efforts to characterize the HRF have focused on measuring the transient, task-evoked BOLD responses to known events or stimuli (i.e. event-related fMRI), where timing information is accurately known. However, recent reports have shown how identification of the peaks of relatively large-amplitude BOLD signal fluctuations in resting-state data may also be used to reliably estimate HRFs without a stimulus (6, 7). This insight has led to the derivation and characterization of the HRF along the entire cortex in resting-state data (8), and the potential use of the HRF as a biomarker to study the effects of development, aging, or pathology (9-11).

Most measurements of the HRF have focused on gray matter (GM), as BOLD effects in white matter (WM) have been reported relatively rarely (12), and often are regressed out as nuisance covariates. Blood flow and volume in WM are only about 25% as large an in GM (13, 14), and the energy requirements for WM functions are usually considered low compared to the cortex (15), so large BOLD effects are not expected. However, there have been several recent reports of successful detection and analyses of WM BOLD signals in both a resting state and after a task (16, 17), and these have led to increased awareness of the relationships between GM activity and WM BOLD signals, and of the correlations in BOLD signals between different WM and GM regions in a resting state (18, 19). Given the different composition, vasculature and functions of GM and WM, their energy use and neurovascular coupling may be different. Characterizing the HRF in WM accurately is relevant for detection, quantification and interpretation of BOLD effects. While neural signaling processes demand substantial energy consumption in GM (13, 14), the metabolic support of other processes are believed to dominate the energy budget in WM, in contrast to GM demands (15). These non-signaling metabolic requirements maintain resting potentials on cell membranes and support general housekeeping including the maintenance of myelin, and are a feature of energy-use in non-neuronal glial cells. Thus, BOLD signal fluctuations in WM may be driven by different energy demands and cell/tissue types than in GM, and the HRF may reflect differences in neurovascular coupling. Recent reports have confirmed that WM HRFs in task-based fMRI are similar to but different from those in GM (16, 17).

We aimed to provide a comprehensive characterization of the HRF in WM across a population, including accurate measurements of its shape and its variation along and between WM pathways, using resting-state fMRI acquisitions. We also for the first time relate features of the HRF to microstructural and physiological properties of tissue derived from diffusion MRI. The relationships of HRFs between tissue types, and the measured differences in the HRF along and between WM pathways, suggest there are different energy requirements between different WM pathways, and the results are consistent with the hypothesis that the HRF in WM reflects the metabolic needs of different tissue components than those seen in GM.

## Methods

### Data

The data and HRF estimates closely followed the approach of Li et al (2021). (20). 199 subjects were randomly selected from the HCP S1200 release (87 M/112 F; age 22-35). The images included resting-state fMRI, T1 weighted MRI, and diffusion MRI. The imaging protocols have been described in detail in previous reports (21). Briefly, data were acquired using a 3T Siemens Skyra scanner (Siemens AG, Erlanger, Germany). The resting-state data were acquired using multiband gradient-echo echo-planar imaging (EPI). Each session consisted of two runs (with left-to-right and right-to-left phase encoding) of 14 min and 33 s each (TR = 720 ms, TE = 33.1 ms, voxel size = 2 mm isotropic, number of volumes = 1200). Physiological data, including cardiac and respiratory signals, were recorded during fMRI acquisitions. The diffusion MRI were acquired using a multiband spin-echo EPI, again with right-to-left and left-to-right phase encoding polarities (TR=5520 ms, TE =89.5 s, voxel size =1.25 mm isotropic, 3 shells of b=1000, 2000, and 3000 s/mm2). T1-weighted images were acquired using a 3D magnetization-prepared rapid acquisition with gradient echo (MPRAGE) sequence. (TR = 2400 ms, TE = 2.14 s, voxels size = 0.7 mm isotropic).

### Preprocessing

Images were pre-processed through the minimal pre-processing (MPP) pipelines (22) of the HCP. T1-weighted images were non-linearly registered to MNI space using FNIRT (23) and subsequently Freesurfer produced surface and volume parcellations as well as morphometric measurements (24). For fMRI, the analysis pipeline included motion correction, distortion correction using reversed-phase encoding directions, and non-linear registration to MNI space. We performed additional processing including regression of nuisance variables, including head movement parameters (using one of the outputs of motion correction in the MPP pipeline), and cardiac and respiratory noise modeled by the RETROICOR approach (25), and followed by a correction for linear trends and temporal filtering with a band-pass filter (0.01 – 0.08 Hz). A group-wise WM mask was reconstructed by averaging the WM parcellations that were derived from Freesurfer across all subjects and thresholded at 0.9. Then the fMRI data were spatially smoothed within the WM mask with a 4 mm FWHM Gaussian kernel. Similarly, data were also smoothed within a GM mask that was reconstructed in a similar manner but using a lower threshold (0.6) due to higher individual variabilities in GM. For diffusion images, the MPP pipeline included a zero-gradient intensity normalization, EPI distortion correction using reversed-phase encoding directions, and, again, non-linear registration to MNI space.

#### HRF Estimation

HRFs were estimated from resting-state time courses in each subject using a blind deconvolution approach (6, 11) implemented using the rsHRF toolbox (8). The method requires no prior hypothesis about the HRF and is based on the notion that relatively large amplitude BOLD signal peaks represent the occurrence of separable, major, spontaneous events. In our study, such events were detected as peaks beyond a specified threshold (here, greater than 1.5 standard deviations over the mean). For each event, a general linear model was fitted using a linear combination of two double gamma functions together with a temporal derivative to fit the derived waveforms. The double gamma functions together with temporal derivative are capable of modeling an initial dip, a time delay and a later undershoot in the response (26, 27).

### HRF Features

After the HRF had been estimated for each voxel in MNI space, features of the HRF were extracted and visualized as separate parametric maps as shown in **Figure 1**. These included (1) the full width at half maximum (FWHM), a measure of BOLD response duration in seconds; (2) the peak height (Height), a measure of maximum normalized signal response; (3) the Time to Peak, a measure of response latency in seconds; (4) the Time to Dip (where evident), the time to reduce from baseline to the most negative early response; (5) the Dip Height (where evident), corresponding to the maximum normalized signal decrease from baseline (recorded as a negative value); (6) the Peak Integral, the area under the curve of the positive BOLD response; and (7) the Dip Integral, the area under the curve of the early negative BOLD response.

**Figure 1.**
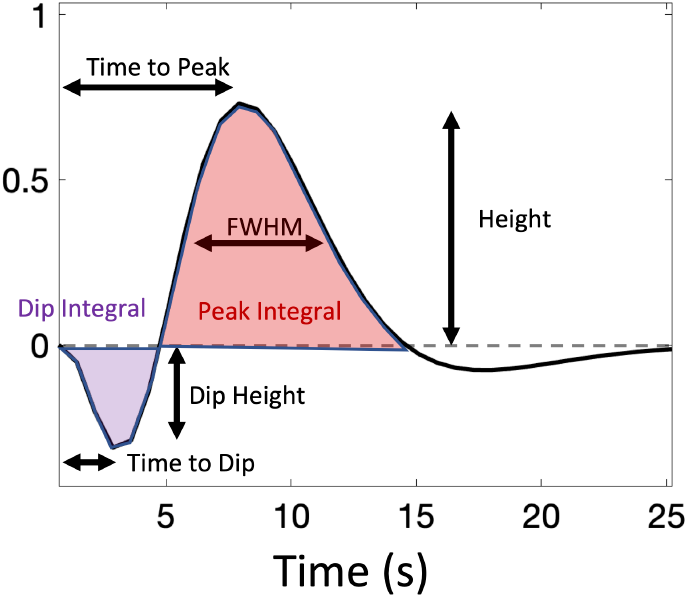
The Hemodynamic response function (HRF) and its features: Time to Peak, Height, Full-width Half-Max (FWHM), Peak Integral, Time to Dip, Dip Height, and Dip Integral.

### Microstructure Features

Diffusion MRI scans were used to derive approximate measures of average tissue microstructure, using the Diffusion Tensor Imaging (DTI) technique and the Neurite Orientation Dispersion and Density Imaging (NODDI) technique (28).

DTI characterizes the magnitude, degree of anisotropy, and orientation of directional diffusion. From this, measures of fractional anisotropy (FA), mean diffusivity (MD), axial diffusivity (AD), and radial diffusivity were derived. The measures are sensitive to a variety of microstructural features. For example, FA may reflect coherence of membranes and restrictions, whereas RD is sensitive to myelination and axonal number and packing density (29, 30). The NODDI model represents the signal in each voxel as the sum of three tissue compartments – intra-neurite (sometimes called intra-cellular), extra-neurite, and cerebral spinal fluid (CSF). The intra-neurite compartment is composed of neurites (modelled as zero-radius sticks) with a distribution of directions that includes both an average direction and a spread of orientations around that direction. Thus, application of NODDI produced a set of parametric maps, averaged across the population, of (1) an isotropic volume fraction (ISOVF), (2) the intra-neurite volume fraction (or neurite density index; NDI), and (3) an orientation dispersion index (OD) where a higher value represents a larger spread of axon orientations. NODDI fitting on each voxel and subject was performed using the accelerated microstructure imaging via complex optimization (AMICO) method (31).

Of particular importance, the quantity (1-NDI) represents the relaxation weighted volume fraction of all non-neurite components, which includes anything that does not display stick-like diffusion signal properties such as glial cells and some extracellular spaces.

### Along-tract quantification

To investigate HRF features of specific WM pathways, we analyzed a set of expertly delineated bundles in MNI space (32). For this work, we focused on 15 major association, projection, and commissural pathways of the brain: the arcuate fasciculus (AF, left and right), corticospinal tract (CST, left and right), inferior longitudinal fasciculus (ILF, left and right), superior longitudinal fasciculus (SLF, left and right), optic radiation (OR, left and right), frontal aslant tract (FAT, left and. right), uncinate fasciculus (UF, left and right), and the corpus callosum.

HRF features were quantified along each major WM pathway using the along-fiber quantification technique on the population averaged WM tracts defined by Yeatman et al. (2012). (33). Each pathway was segmented into N=20 points. Along-pathway mean and standard deviations of each HRF feature were derived for each position along the pathway.

#### Statistical analysis

Linear mixed-effects modeling was used to ask whether HRF features change along pathways: the regression equation used for each feature, y, was;

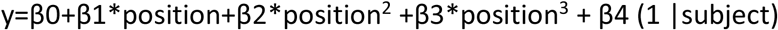

where subject was considered as a random effect (i.e., allowing for a subject-specific intercept), and we tested the null hypothesis that all fixed-effect coefficients equal zero (β1= β2= β3=0) using an F-test. A rejection of the null hypothesis suggests that a feature, y, changes along a pathway. We additionally performed a one-way analysis of variance (ANOVA) to test whether there were significant differences between pathways. We also evaluated whether the variation of each feature is different along each pathway, where the variation is defined as the ratio of the minimum to the maximum value of the feature along the pathway.

## Results

The results provide information in response to several questions we sought to address.

### [a] What is the average HRF across a population?

**Figure 2** shows WM and GM masks, and parametric maps of each of the 7 features of the HRF averaged across the population. WM and GM HRFs are qualitatively different, and all features show significant contrast between the tissues.

**Figure 2.**
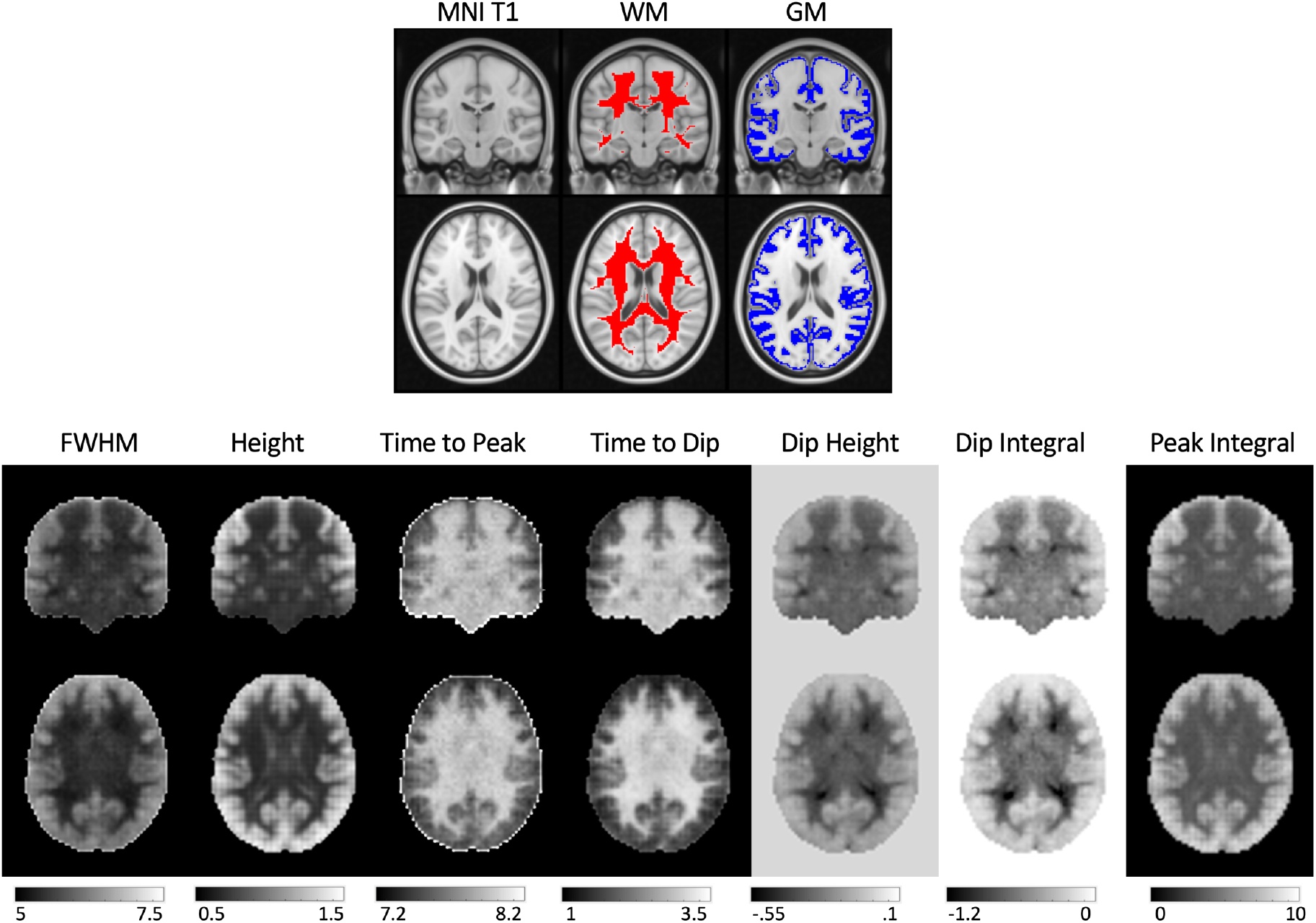
Population maps of HRF features show qualitative differences between GM and WM. (Top) Shown are the MNI T1, WM and GM masks for anatomical reference. (Bottom) Population-averaged features of the HRF are shown for FWHM, Height, PSC, time to Peak, Time to Dip, and Dip Height.

**Figure 3** quantifies differences in each of the features of the HRFs between WM and GM, and the distributions of values within each tissue. The GM HRF shows a typical canonical response, with a barely evident initial dip, whereas the WM shows a consistent and larger initial Dip Height, a smaller Peak Height, and smaller negative undershoot. In general, WM HRFs have a shorter FWHM, lower Peak Height, longer Time to Peak, and larger negative Dip Area, in agreement with previous literature (8). In summary, the HRF shows significant differences between WM and GM tissue types.

**Figure 3.**
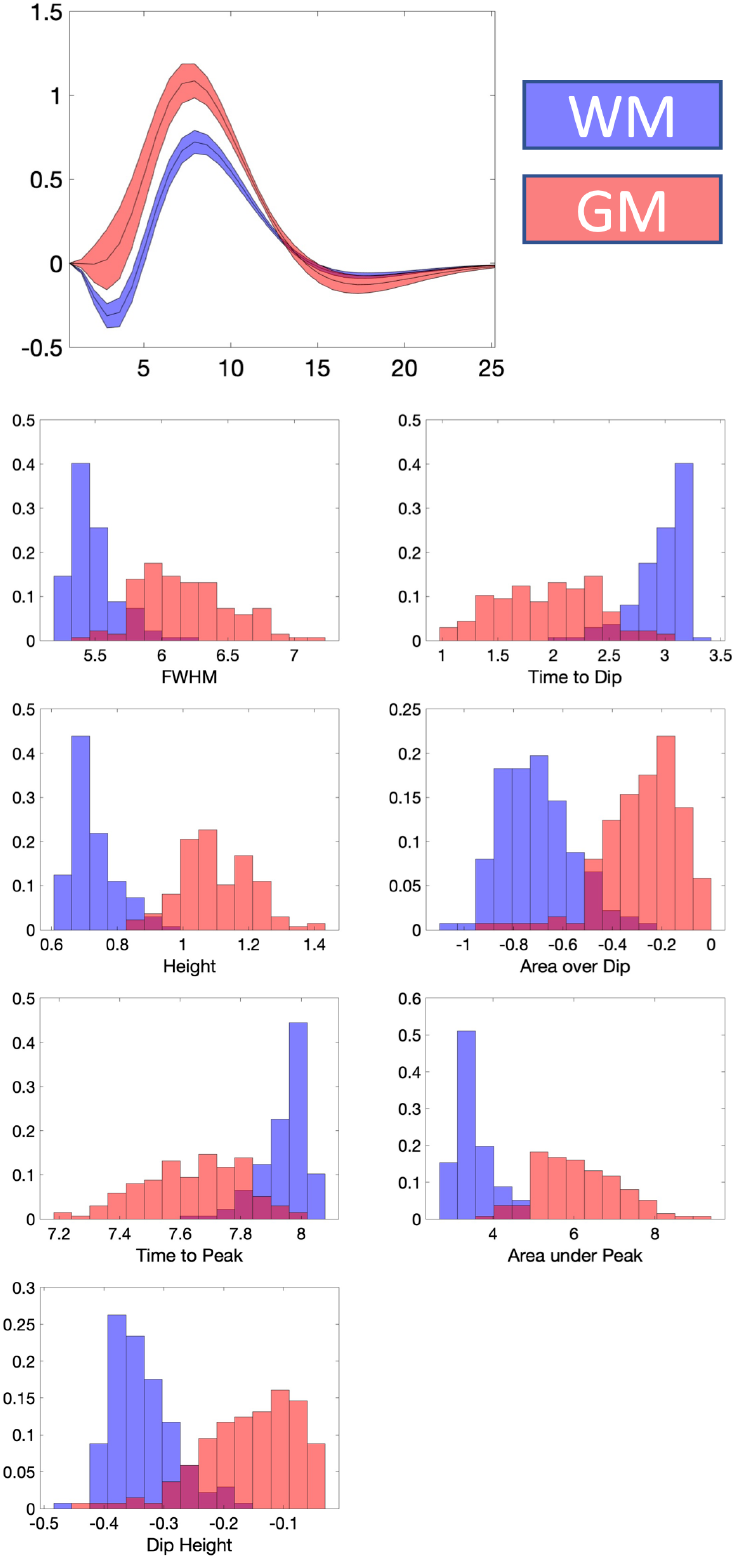
WM and GM tissues have different HRFs and derived features. The population-averaged HRF is shown for WM and GM, displayed as mean and standard deviation across subjects. The distribution of HRF features in WM and GM is displayed as bar plots.

### (b) What are the relationships between HRF features?

**Figure 4** plots the relationships between each pair of HRF features for all voxels in the population-averaged data, with WM voxels plotted as purple and GM as dark-orange. Nearly all features show strong correlations with others, and GM and WM exhibit similar trends but with some notable differences. Examples include strong positive relationships between time to peak and time to dip, dip height and dip integral, but with different slopes for GM and WM. In GM a lower Dip Integral corresponds to a larger, faster positive BOLD increase (larger Height, FWHM, shorter Time to Peak) suggesting areas with a faster flow response to each event. Conversely, WM clearly exhibits larger Dip Integrals overall and the subsequent positive response shows little variation in Height or Time to Peak. In WM there is little covariation of Peak Integral with FWHM, Peak Height or Time to Peak, whereas in GM the Peak Integral has strong positive correlation with the FWHM and Height and negative relationship with Time to Peak.

**Figure 4.**
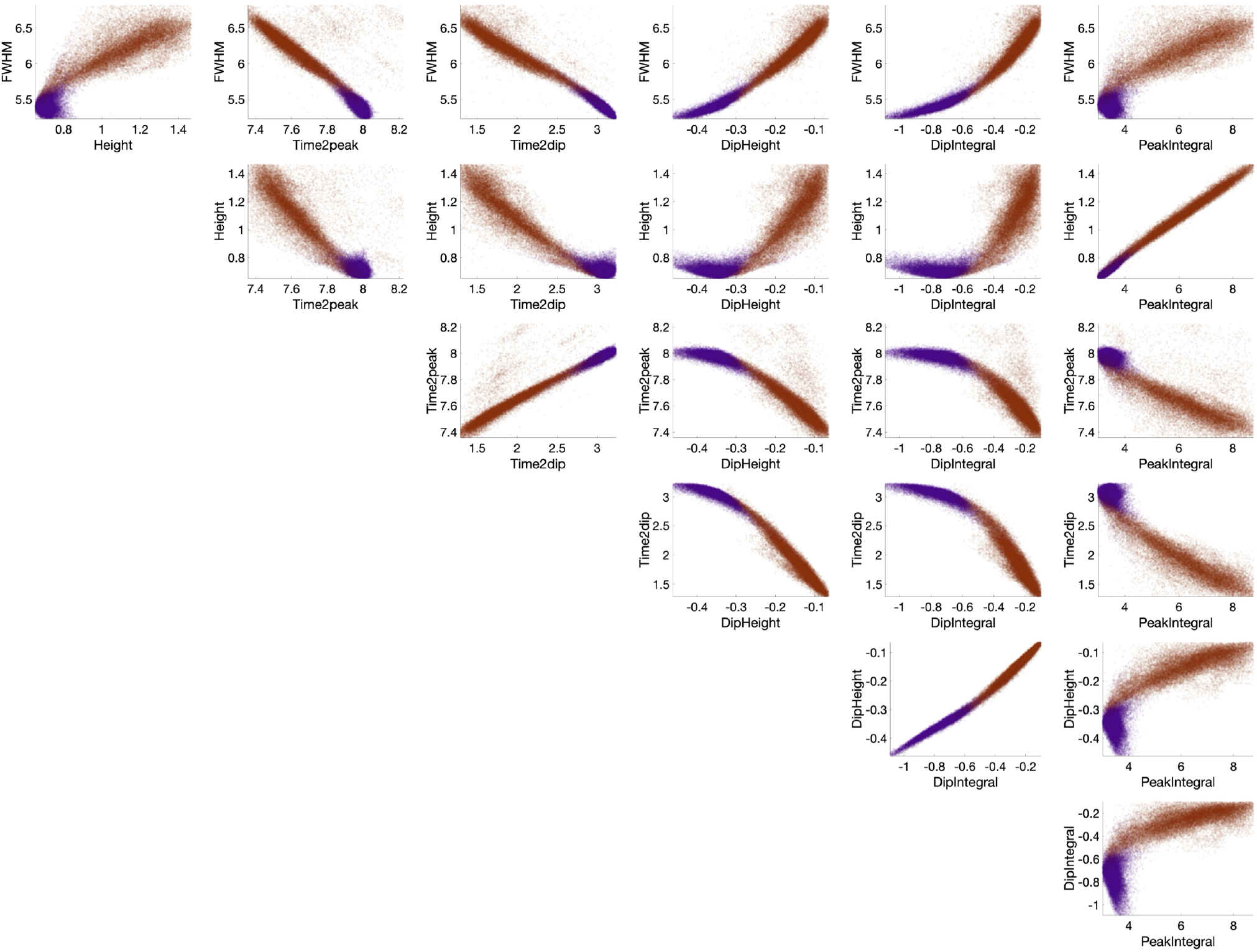
Features of the HRF show strong relationships with each other in both WM and GM. Here, each HRF features is plotted against all other HRF features for all voxels, with WM voxels shown as purple and GM as dark orange.

### (iii) What are the relationships between HRF and microstructure?

**Figure 5** plots the relationships between HRF features and diffusion-derived microstructural features for all voxels in the population-averaged maps. Again, WM and GM are shown as purple and dark orange, respectively. As expected, white matter exhibits different tissue microstructure than GM, with a higher FA, a higher NDI, and lower RD. More interesting, WM and GM show very different relationships between HRF and microstructure. For example, in WM, the area under the negative dip (Dip Integral) decreases (smaller negative value) with increasing NDI and with increasing FA. However, this dip integral shows the opposite trend in GM, decreasing with increasing NDI and increasing FA. Likewise, the FWHM covaries with NDI and RD but with differing relationships between WM and GM.

**Figure 5.**
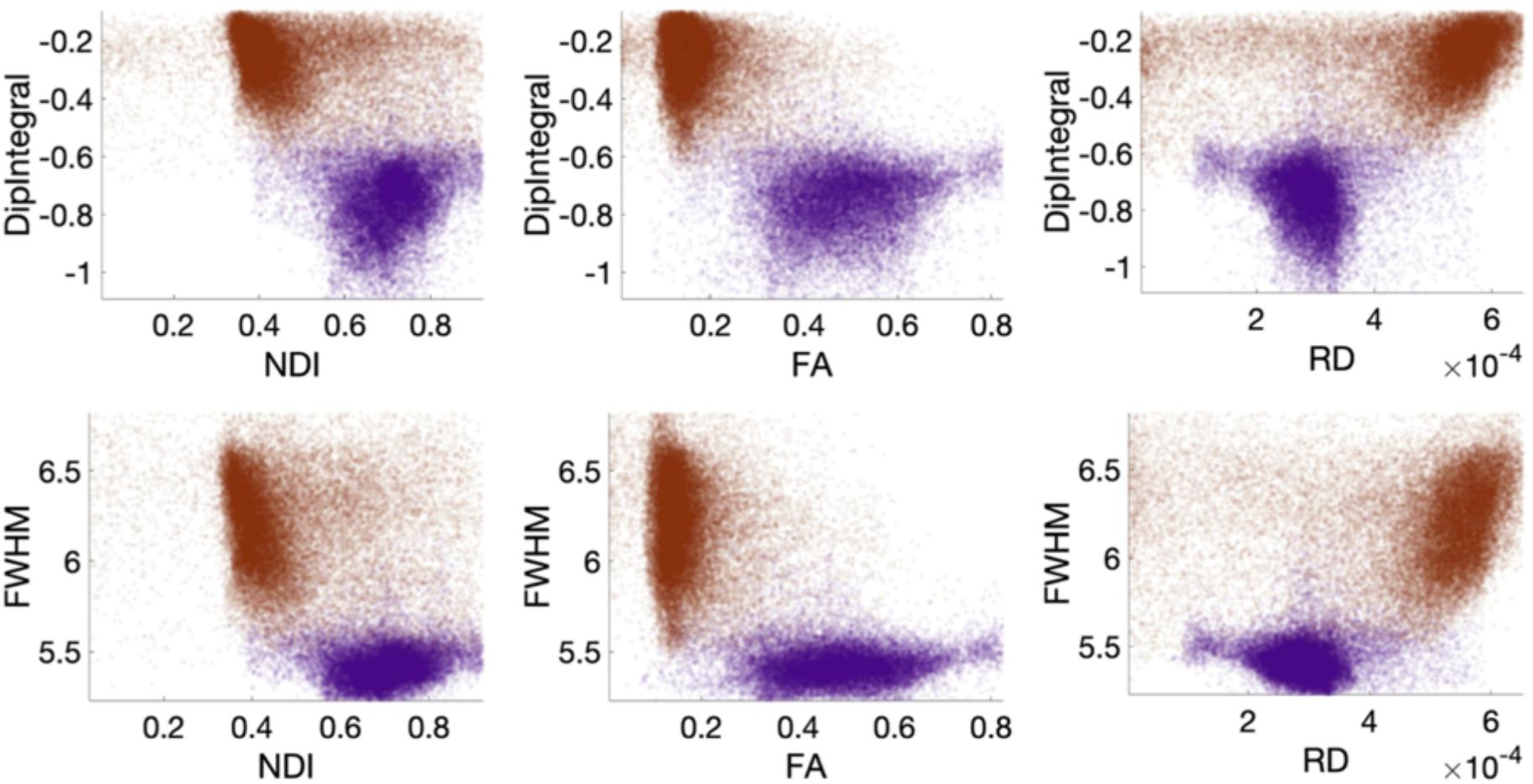
Features of the HRF show distinct relationships with tissue microstructure in WM and GM. Here, HRF features are plotted against microstructure features for all voxels, with WM voxels shown as purple and GM as dark orange.

### (iv) How does the HRF vary along WM pathways?

The HRFs for 15 WM pathways are shown in **Figure 6**. Here, streamlines are color coded from blue to red, with corresponding color-matched HRFs shown averaged across the population. While the overall shape is similar within and across pathways, larger variations are observed near the WM/GM boundary at the ends of the pathway, with smooth trends along pathways. Visually, the peak heights decrease in the core of each WM pathway, whereas, the dip height increases. Pathways such as the ILF, SLF, and OR show visually higher variation across the population.

**Figure 6.**
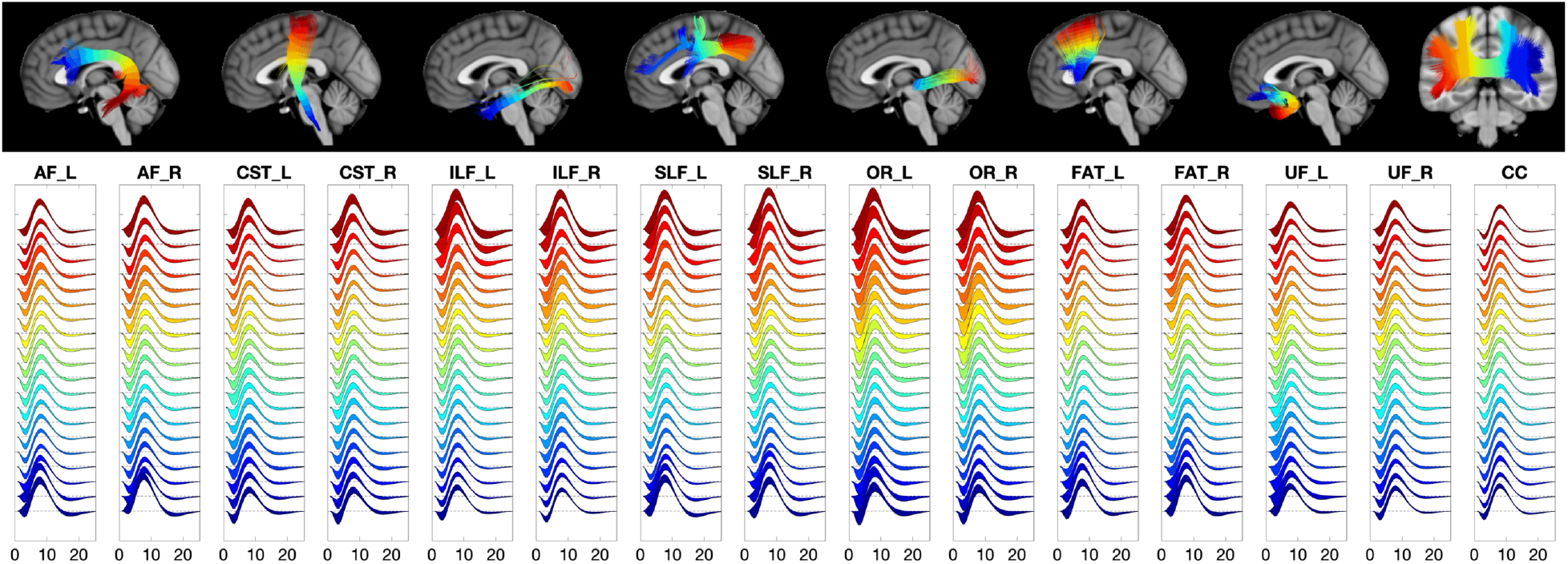
The HRFs are qualitatively different across and along pathways. Visualization of the HRFs along 15 white matter pathways, color-coded from beginning (blue) to end (red) of pathways. Plots show the means and standard deviations across the population.

**Figure 7** shows two exemplar pathways (the arcuate fasciculus and optic radiations) and the corresponding change in HRF features along the pathway. Trends are apparent along pathways, including a decreased height, decreased FWHM, increased time to peak and time to dip, and larger negative dip in the middle core of the pathway. Statistical analysis confirms that all features, of all pathways, significantly change along the pathway from start to end. All p-values for the F-tests are given as supplementary information.

**Figure 7.**
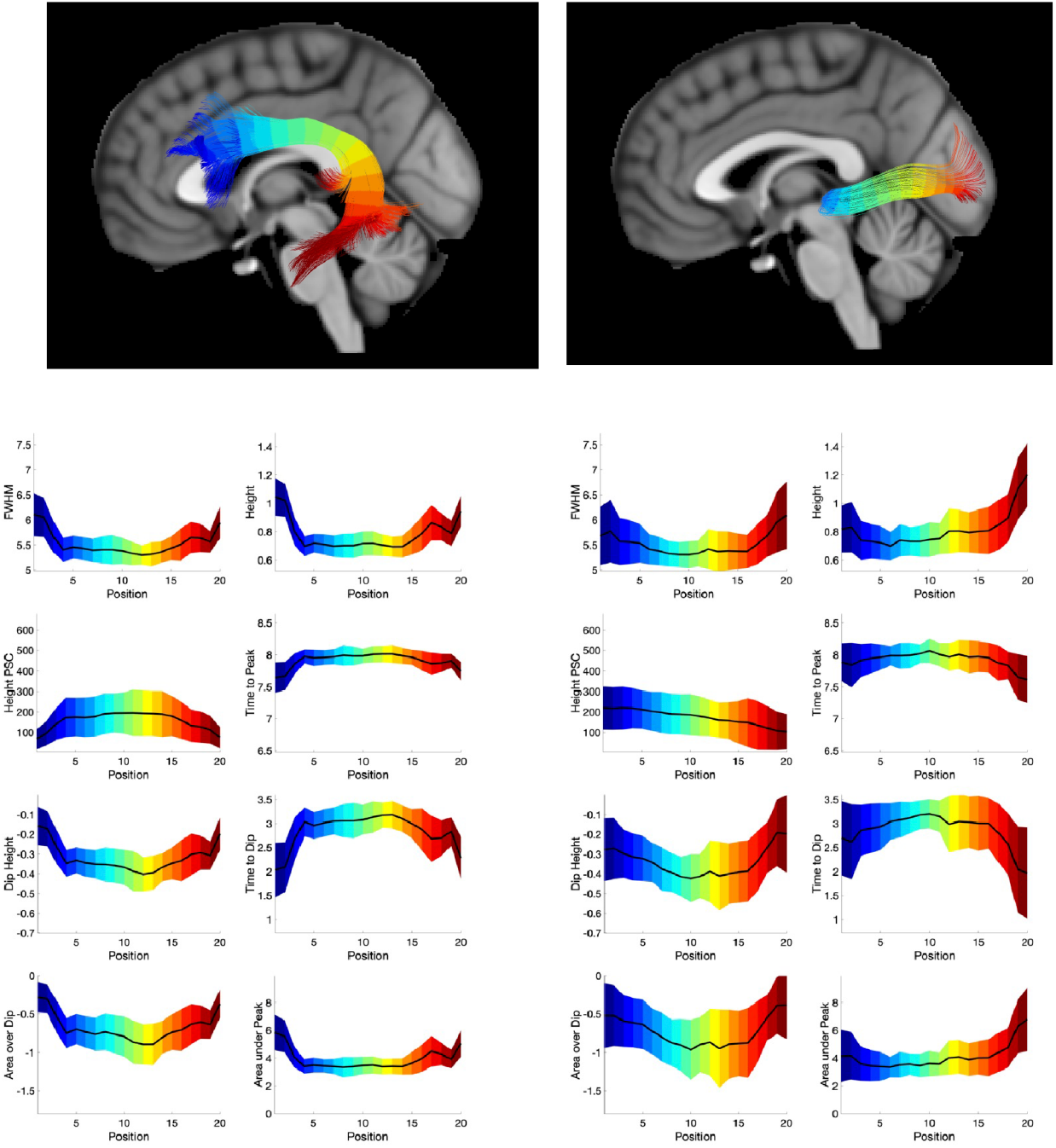
HRFs change along pathways. Exemplar pathways of the arcuate fasciculus and optic radiations are shown, with the 8 HRF features and their changes along the pathway. Color-coded to match the visualizations from beginning (blue) to end (red) of pathway. All features statistically significantly changes along pathways.

### (v) How does the HRF vary between WM pathways?

We aimed to extract a measure of changes in each HRF feature along pathways by taking the ratio of the range of that feature over the maximum value, calling this “% Variation”. The % Variations of each feature, for all features along every pathway, are shown in **Figure. 8**. We find that several pathways show more variation in HRF features than others, including the optic radiations, and arcuate fasciculi, and that the observed variations are significantly different across all pathways.

**Figure 8.**
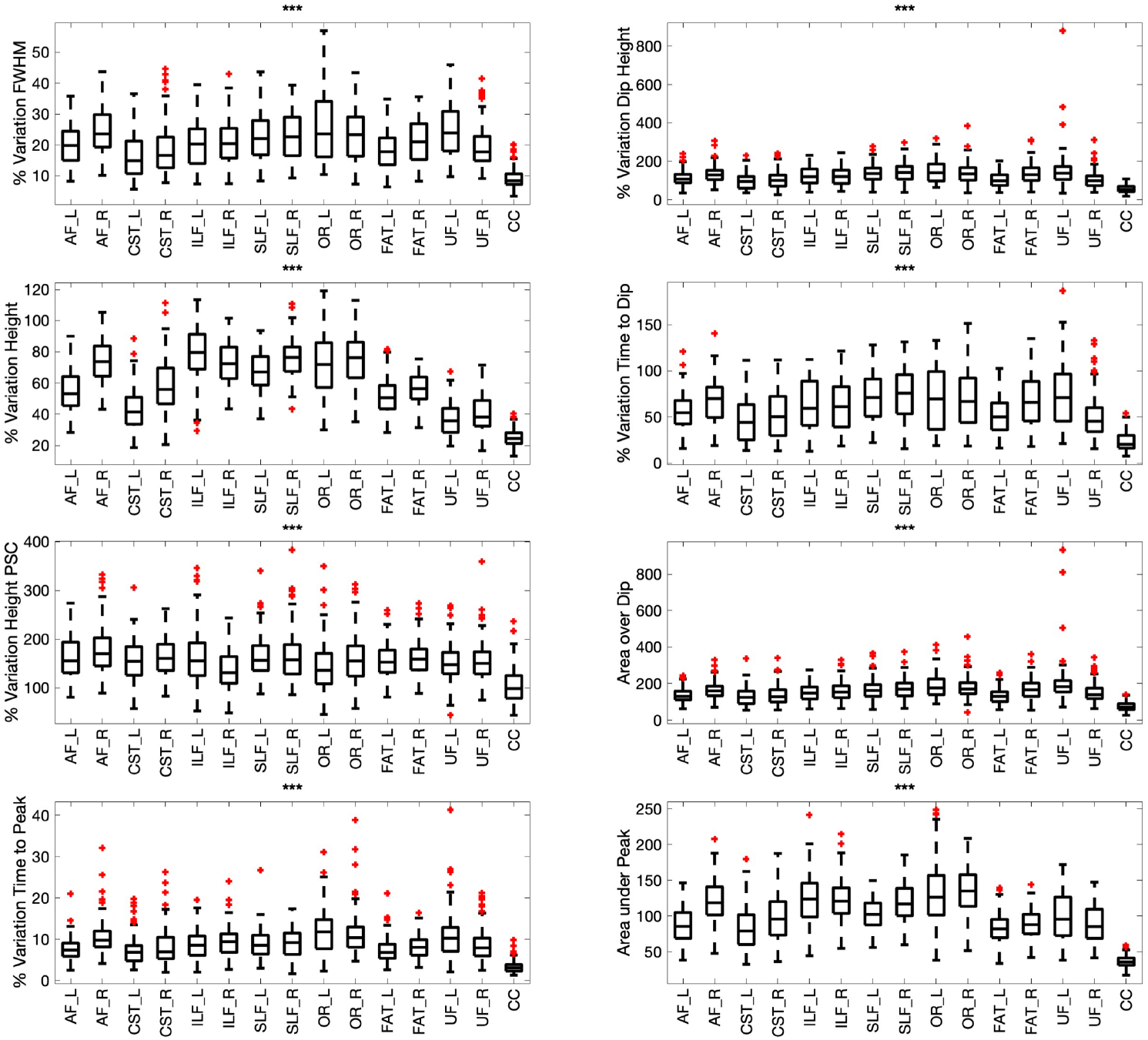
The variation along pathways (for each feature) is quantified by taking the range (max – min) divided by the maximum value to indicate the “% variation” along pathways. This variation is shown for all pathways, and is statistically significantly different between pathways.

## Discussion

We have quantified variations of the HRF of resting state fMRI within and across white matter pathways and cortical grey matter by creating population-averaged parametric maps of HRFs and their derived features. We find that, following transient fluctuations in resting-state signal, WM BOLD responses have smaller peaks and are slower to change, than GM responses, and also feature a prominent negative initial dip. Second, we find strong links between different features of the HRF, with similar patterns observed in both WM and GM, suggesting similar factors affect the dynamics and evolution of the response function. However, the tissue microstructural environment differs significantly between these tissue types, and the dynamics of the HRF show different dependencies on the underlying structure. Next, we find that features of the HRF change significantly along white matter pathways, and are much different in the deep WM than at the superficial WM. The variation observed along pathways also differs between pathways. Together, this suggests that, much like in GM, changes in flow and/or oxygenation related to variations in baseline conditions are different for different parts of the WM. These differences in HRF features may be relevant for understanding the biophysical basis of BOLD effects in WM.

### Biophysical basis of BOLD signal in WM

The origins of BOLD effects in WM are not well understood though they share many features with BOLD effects in GM and thus a common explanation of their biophysical basis is plausible. The HRF features include an initial negative dip, indicating an increase in the concentration of deoxyhemoglobin within tissue, followed by a positive peak that extends for several seconds, indicating an increase in blood flow and volume, a net decrease or washout of deoxyhemoglobin, and then a return to equilibrium. The times for flow to increase and subsequently return to baseline are likely determined by the overall density and dimensions (diameters and spacings) of the vasculature involved (34-36). There is converging evidence that BOLD effects in WM are coupled with changes in neural activity within GM (12, 37, 38) but it is not proven whether this reflects an intrinsic metabolic demand in WM or potentially flow effects from adjacent GM, though the latter seems less likely given the nature of the vasculature within WM and the extent of the BOLD effects detected.

As pointed out by several previous reports (15, 39), the metabolic demands of neural processes in WM and GM are different and only a relatively small fraction of the energy budget is WM is believed to be required for signal transmission along axons. Thus, whereas in GM the increase in oxygen consumption is directly related to intrinsic neural activity, in WM the BOLD responses to activity within the cortex are likely triggered by a different requirement. Potential sources of increased metabolic demand include the glial and other non-neuronal cells that constitute a large fraction of WM (40). The two macroglia oligodendrocytes and astrocytes are the most abundant cell types. In addition to their roles in maintaining microstructure and production of essential lipids, glial cells are involved in a host of processes associated with brain function including the regulation of ionic balances, pH, neurotransmitter actions and other requirements. We postulate that BOLD signals in WM are related to the metabolic requirements of the glial cells and thus predict there should be correlations between glial content and features of the HRF.

To investigate this potential connection, we produced atlases of microstructure features derived from diffusion MRI. From the diffusion models it is possible to derive voxel-wise maps of parameters including FA and NDI. FA is highly sensitive to the fractional composition of tissue that is myelinated neurons. Complementing the neuronal fractional volume, the extracellular volume fraction (ECVF=1-NDI) is a measure of the volume fraction within a voxel that is not neuronal and thus is a surrogate metric of the glial cell volume fraction. Figure 5 shows the correlation between ECVF and the negative dip of the HRF for WM and GM. The area under the negative dip of the BOLD HRF is presumed to indicate a larger metabolic demand within the tissue that causes tissue pO2 to decrease transiently. In WM, this correlates positively with the ECVF and negatively with the FA. On the other hand, it correlates negatively with the ECVF in GM, consistent with the negative dip being smaller for cortex containing a small neuronal volume fraction.

### Variation along and between pathways

The use of diffusion tractography and along-fiber quantification (33) has proven useful in identifying location in the WM, and determining what changes occur, and where they occur, in development, disorders, and disease. With the introduction of the rsHRF toolbox used here, Wu and colleagues noted differences in WM and GM HRF’s (8), and found areas in the WM that showed HRF alterations in schizophrenia and attention-deficit hyperactivity disorder (ADHD). Here, we show that along-tract profiling is possible with non-diffusion-based measures, and offers the ability to add specificity to identify which pathways may be affected and where along that pathway experiences alterations.

In addition to showing feasibility of along-tract profiling of HRF features, our results also suggest that energy demands are different across different pathways in a resting state. This parallels the existence of resting-state networks in the cortex. Just as various networks may be characterized for aspects of attention, memory, motor, sensory systems, the white matter fibers are structures often associated with unique functional roles, so it is unsurprising that they may require different energy processes on different time-scales. In fact, recent work (19) shows that by using appropriate analysis, the white matter may be robustly parcellated into functional networks. The use of *a priori* white matter pathways, as opposed to these data-driven approaches, allows us to associate changes in blood flow and oxygenation to specific white matter pathways with well-defined functional roles (41).

There are several directions to pursue to extend this research. First, the use of the HRF as a potential biomarker would be strengthened if WM HRFs were shown to be changed in altered states/disorders as shown in GM (42-44), and if results are congruent with those suggesting deterioration of specific structural systems as studied with diffusion tractography (45, 46). Studies like this would facilitate comprehensive evaluation and integration of structure and function. Second, HRFs can be quantified in task-based fMRI experiments. Experiments could be designed to elicit responses to study specific structure-function relationships, or for brain-wide activation (47) that may facilitate normative pathway-specific responses during the signaling process. Finally, because pathways are known to spatially overlap in the brain, creating ‘crossing-fiber’ (48, 49) and ‘bottleneck’ (50, 51) problems in the tractography process, it may be possible to disentangle multiple responses within the same voxel through a deconvolution process, or by using microstructure-informed or tractography-informed filtering (31, 52-54), in order to extract multiple HRFs. within the same WM voxel that could be analyzed on a fiber-element basis (55).

## Conclusion

We have characterized the HRF of resting state fMRI within and across white matter pathways by creating population-averaged maps of the HRF and features of the HRF, and show that the HRF shows significant differences between WM and GM tissue types, features of the HRF show strong relationships with both other HRF features and with microstructural features, and that these trends differ between tissue types. We further find that the HRF varies significantly along pathways, and is different between different WM pathways. Together, this suggests that, much like in GM, changes in flow and/or oxygenation related to variations in baseline conditions are different for different parts of the WM. Moreover, we provide evidence suggesting the BOLD responses in WM reflect tissue components different from myelinated axons. These differences in HRF features may be relevant for understanding the biophysical basis of BOLD effects in WM.

## Acknowledgements

This work was supported by the National Institutes of Health (NIH) grants K01EB032898, R01 NS093669, 5R01EB017230 and R01 NS113832, and Vanderbilt Discovery Grant FF600670. Imaging data were provided by the Human Connectome Project, WU-Minn Consortium (Principal Investigators: David Van Essen and Kamil Ugurbil; 1U54MH091657) funded by the 16 NIH Institutes and Centers that support the NIH Blueprint for Neuroscience Research; and by the McDonnell Center for Systems Neuroscience at Washington University.

## Notes

### Competing Interest Statement

The authors have declared no competing interest.

